# Within and across-trial dynamics of human EEG reveal cooperative interplay between reinforcement learning and working memory

**DOI:** 10.1101/184812

**Authors:** Anne GE Collins, Michael J Frank

## Abstract

Learning from rewards and punishments is essential to survival, and facilitates flexible human behavior. It is widely appreciated that multiple cognitive and reinforcement learning systems contribute to behavior, but the nature of their interactions is elusive. Here, we leverage novel methods for extracting trial-by-trial indices of reinforcement learning (RL) and working memory (WM) in human electroencephalography to reveal single trial computations beyond that afforded by behavior alone. Within-trial dynamics confirmed that increases in neural expectation were predictive of reduced neural surprise in the following feedback period, supporting central tenets of RL models. Cross-trial dynamics revealed a cooperative interplay between systems for learning, in which WM contributes expectations to guide RL, despite competition between systems during choice. Together, these results provide a deeper understanding of how multiple neural systems interact for learning and decision making, and facilitate analysis of their disruption in clinical populations.

**One sentence summary:** Decoding of dynamical neural signals in humans reveals cooperation between cognitive and habit learning systems.

## Introduction

When learning a new skill (like driving), humans often rely on explicit instructions indicating how to perform that skill. However, for many problems, these instructions may be too numerous to keep in working memory, and one needs to focus on a subset of them while acquiring large portions of skills by trial and error, or reinforcement learning (RL): “practice makes perfect”. Previous research showed that dual cognitive and incremental RL systems contribute to learning across a range of situations, even when explicit instructions are not provided, and stimulus-response contingencies must be acquired solely by reinforcement (*1*–*7*).

This body of work is motivated by theoretical considerations suggesting that RL and cognitive systems optimize different trade-offs. The RL process statistically integrates reinforcement history to estimate the expected value of choices, in accordance with “model-free” algorithms that guarantee convergence, but are slow and inflexible (*8*). This process is widely thought to be implemented in cortico-basal ganglia loops and their innervation by dopaminergic signals (*9, 10*). In contrast, the cognitive system facilitates more flexible and rapid learning, but is limited by working memory (WM) capacity, is subject to forgetting, and is evidenced by differential efficiency of learning in simple and complex environments (*6*). The WM system is a primitive for more “model-based”, or goal directed cognitive processes and is thought to depend on prefrontal cortex among other regions (*1, 3, 11*).

Although it is well established that multiple systems contribute to learning, their interactions are poorly understood. We recently showed that frontostriatal reward prediction error (RPE) signals – the canonical neuroimaging signature of model-free RL – are more strongly represented (*3*), and behavioral value learning is actually enhanced, under high compared to low WM load (*7*). However, it remains unclear whether these interactions reflect a competitive or cooperative process (see below). Here, we combine computational modeling, electro-encephalography (EEG), and decoding to provide insight into this issue. Specifically, EEG allows us to interrogate within-trial dynamics of the two systems, and how they are combined to converge on a single decision and interpret an outcome. We use computational modeling to quantify variables involved in RL and WM, and decoding to identify their signatures in EEG. First, we confirm that EEG markers of reward expectation at decision onset are negatively coupled with markers of RPE in the subsequent feedback period within the same trial, as predicted by axiomatic tenets of RL, but never directly shown in neural data. Second, we predicted that we would see markers of RL processing earlier than those of WM in the neural signal, given that the latter process is more cognitively costly. Finally, we investigate whether the two systems are purely independent, or if they interact with each other. As noted above, earlier work has hinted that WM and executive functions might interfere with, or modify, RL computations (*3, 4, 7, 12*), but it remains elusive whether these interactions are competitive or cooperative. We leverage model-informed within- and across-trial analyses of EEG decoding signals to arbitrate between these two possibilities. We establish the existence of a cooperative interaction between WM and RL, whereby the RL process is counter-intuitively weakened when the learning environment is least complex (i.e., WM load is lowest).

## Results

To parse out contributions of RL and WM to learning, we used our RLWM task (Figure 1a) (*1*–*3, 7*) while recording EEG (see Methods). Participants learned via reinforcement to select one of three actions for each visual stimulus. WM demands were manipulated by varying across blocks the number of stimuli (*ns* or *set-size*) to be learned from 1 to 6, with similarly varying *delays*.

**Figure 1:**
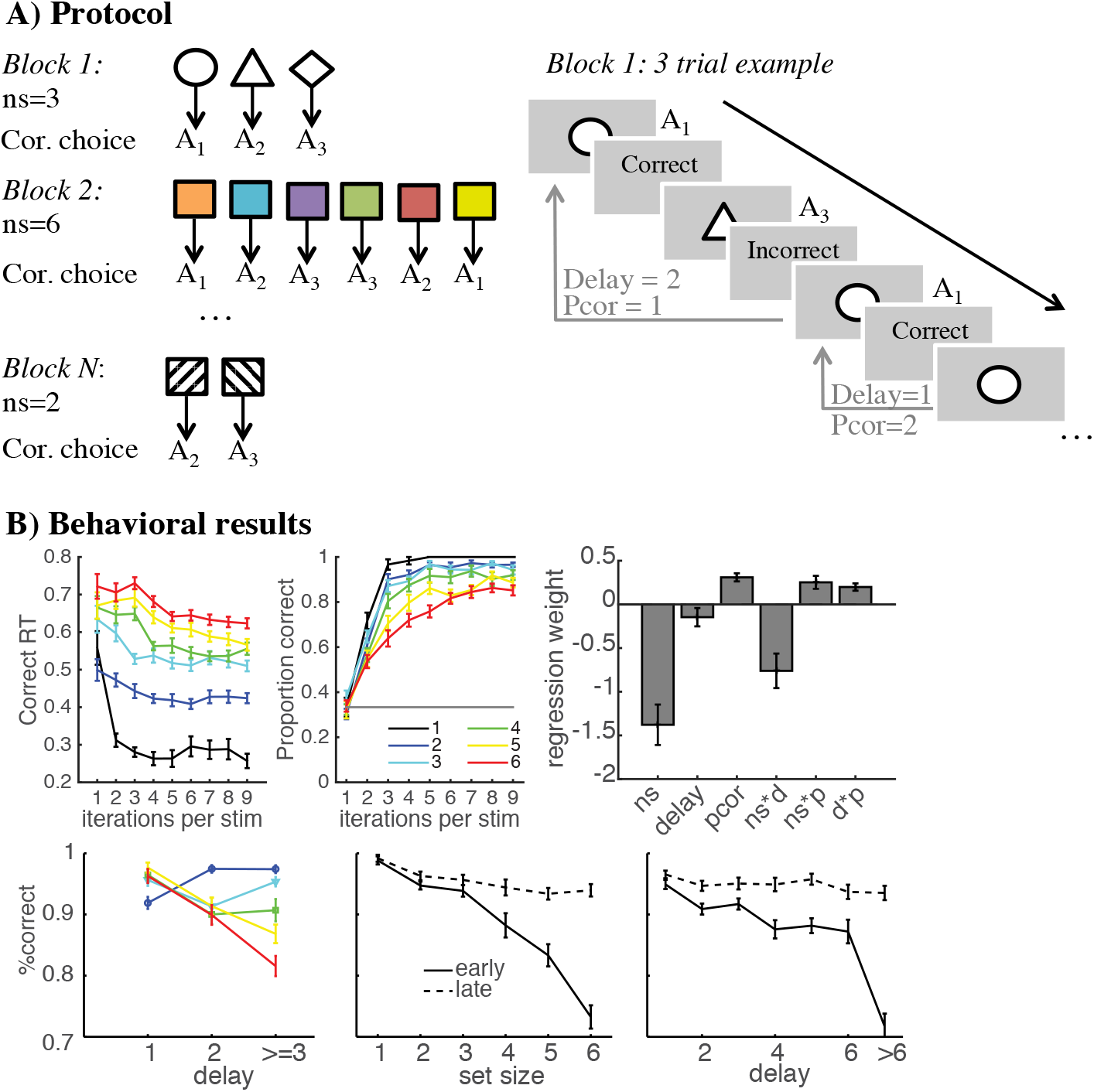
Experimental protocol and behavioral results. **A)** In each block, participants use deterministic reward feedback to learn which of three actions to select for each stimulus image. The set size (or number of stimuli, *ns*) varies from 1 to 6 across blocks. **B)** Top left, middle – reaction times and performance learning curves for each set size as a function of number of iterations of a stimulus. Top right – logistic regression weights show contributions of working memory (smaller set sizes and smaller delays facilitate performance) and reinforcement learning (incremental effects of previous correct trials (*pcor*) for a stimulus), and their interactions. The bottom row shows that these interactions are mediated by greater effects of delay in high set sizes (left), and reduced effects of both set size and delay as learning progresses from early to late in a block (middle and right), suggestive of a transition from WM to RL (*1*).

Behavioral results from 40 participants replicated previous findings implicating separable RL and WM systems, with the relative contribution of WM decreasing with learning. First, participants were more likely to select the correct choice as the number of previous correct (*pcor*) trials accumulated, (Fig. 1B), a basic marker of incremental RL (t(39)=9.1,p<10-4). Second, correct performance was more rapidly attained in lower set sizes and declined with increasing set-sizes (*ns*; t(39)=-5.4, p<10-4) and delays (t(39)=- 3.1;p=.004;), with delay effects amplified under high loads (t=-4.2,p=0.0002), consistent with contributions of a capacity- and maintenance-limited WM system. Finally, interactions between the three factors showed that set-size and delay effects decreased with learning (ts>3.2,p<0.003), confirming a shift from WM to RL with experience (Figure 1B) (*1*–*3, 7*).

### Trial-by-trial decoding of model-based indices of RL and WM

We used our previously developed computational model to quantitatively estimate the contributions of RL and WM to each participant’s behavior. The model includes a standard model-free RL module, which estimates expected “Q” value of stimulus-action pairs and incrementally updates those values on each trial in proportion to the reward prediction error (RPE). This module is complemented by a WM module that assumes perfect memory of the last trial’s stimulus-action-outcome transition, but has limits on both capacity *K* (number of items that can be held in mind, such that the probability of recall *p = K/ns*) and on maintenance (memory for transitions is decayed on each subsequent trial, due to forgetting / updating of intervening items). Model selection confirmed that the RLWM model quantitatively fit participants’ behavior better than other models that assume only a single process (see Fig. 2A), and simulations of the RLWM model captured participants’ patterns of behavior (Fig. 2B-C, Fig. S1). We then extracted trial-by-trial estimates of the expected Q-value and RPE from the RL module (thus factoring out WM contributions to behavior), as a quantity of interest for model-based analysis of EEG.

**Figure 2:**
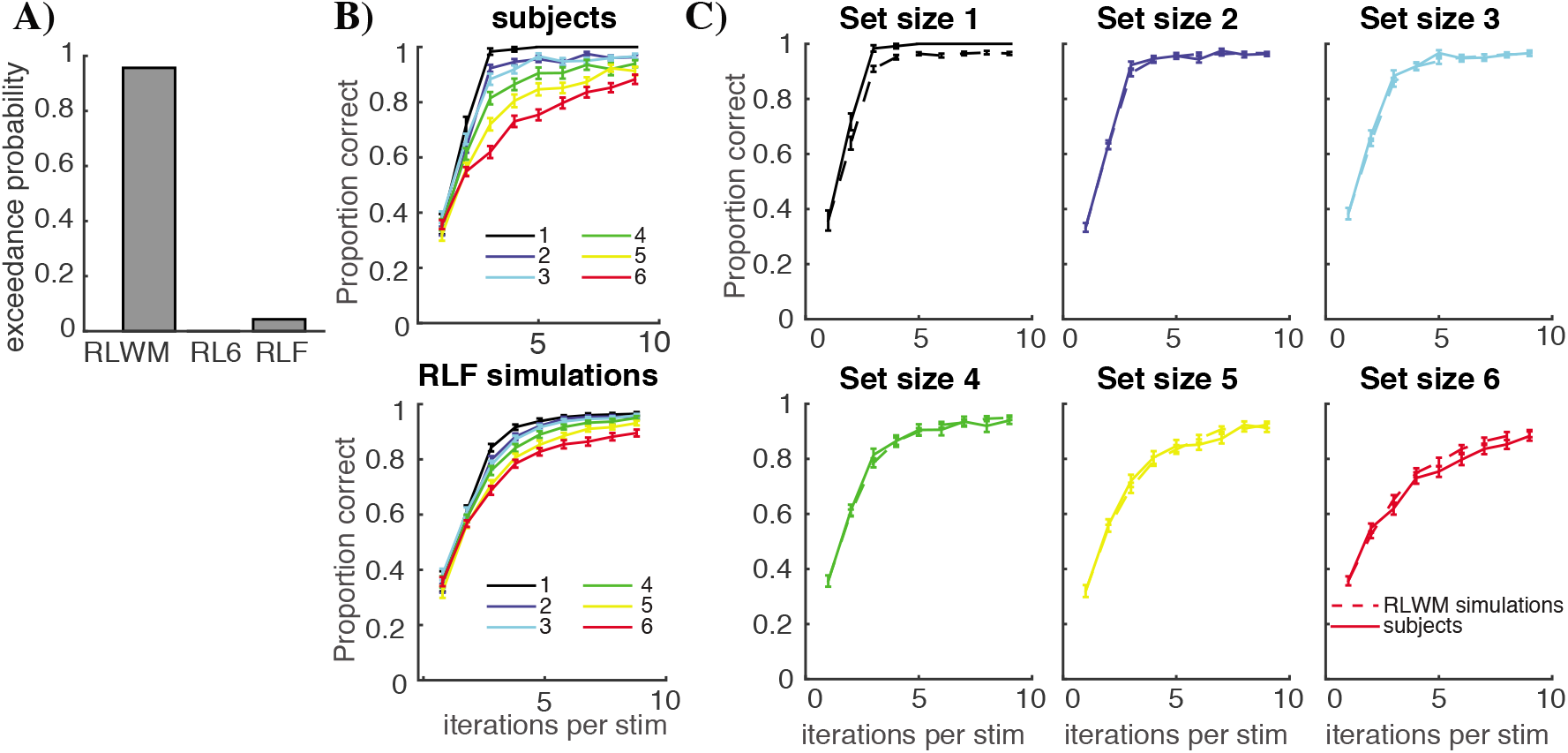
Model validation. **A)** RLWM model provides superior fit to trial-by-trial data compared to competing single actor models, assessed by exceedance probability. RL6 is a standard RL model allowing separate learning rates per set size; RLF is a standard RL model with forgetting. **B)** Simulations of the RLF model with fit parameters do not capture behavior appropriately. **C)** Simulations of the RLWM model with fit parameters captures learning curves in most set sizes. Simulations were run 1000 times per subject.

To investigate the contributions of RL and WM in the neural signals, we leveraged a trial-by-trial decoding approach to analyzing the EEG data (*13*). We used a regression approach to simultaneously extract the effect of multiple variables of interest on the EEG signal at all time-points and electrodes, using correction for multiple comparisons, while controlling for other factors (such as reaction times), and separating out the role of correlated predictors. We identify clusters of electrodes and time points that show significant sensitivity to each predictor. The main predictors of interest were the set size, the delay, and from the model, the expected Q-value (for stimulus-locked analysis) and RPE (for feedback-locked analysis).

In stimulus-locked EEG, this analysis yielded significant and widespread effects of all three main regressors, and similar to behavior, an interaction of set size with delay, indicative of WM (Fig. 3A; Fig. S2, S3). Notably, neural markers of Q-values appeared substantially earlier (starting around 230ms post stimulus onset) than those for set-size (peaking around 600m; Fig. 3A,B,C), supporting the existence of two separable processes sensitive to RL and WM within a trial. Moreover, the early signal modulates the scalp voltage distribution in the same way (Fig. 3C) for increasing Q-values (when the RL system has learned more) and increasing delays (when the WM system is less likely to contain the relevant information), and thus putatively signals the early recruitment of the RL system. For feedback-locked analysis, we observed robust effects of RPE, and RPE-modulated by delay, but only very weak effects of set-size and delays (Fig. 4 and Fig. S4).

**Figure 3:**
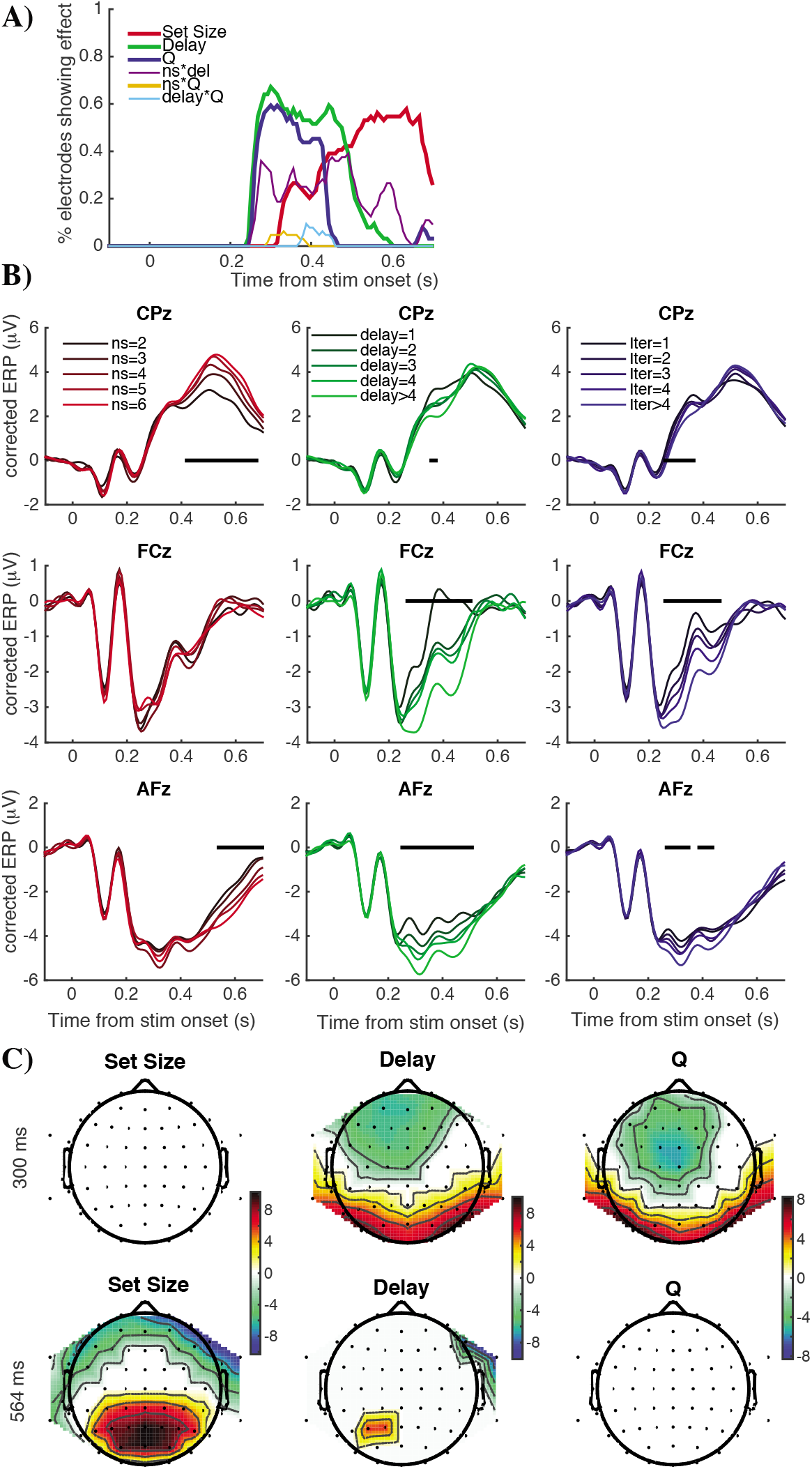
EEG decoding of RL and WM effects during choice. **A)** Proportion of electrodes showing a significant effect (p<0.05 cluster corrected with p<.001 for cluster formation threshold), for each predictor in a multiple regression analysis, against time from stimulus onset. *Q*-values and delays are decoded early (~230ms), whereas set-size *ns* is decoded later (peaking around 600 ms). **B)** Corrected ERPs are plotted to visualize effects accounting for each regression factor (set-size *ns, delay*, stimulus iterations *iter* for three electrodes (CPz, FCz and AFz). **C)** Scalp topography at an early (300ms) and late (564 ms) time point, plotting significance-thresholded average regression weights for the three main predictors. White is non significant; warm colors are positive; cold colors negative.

**Figure 4:**
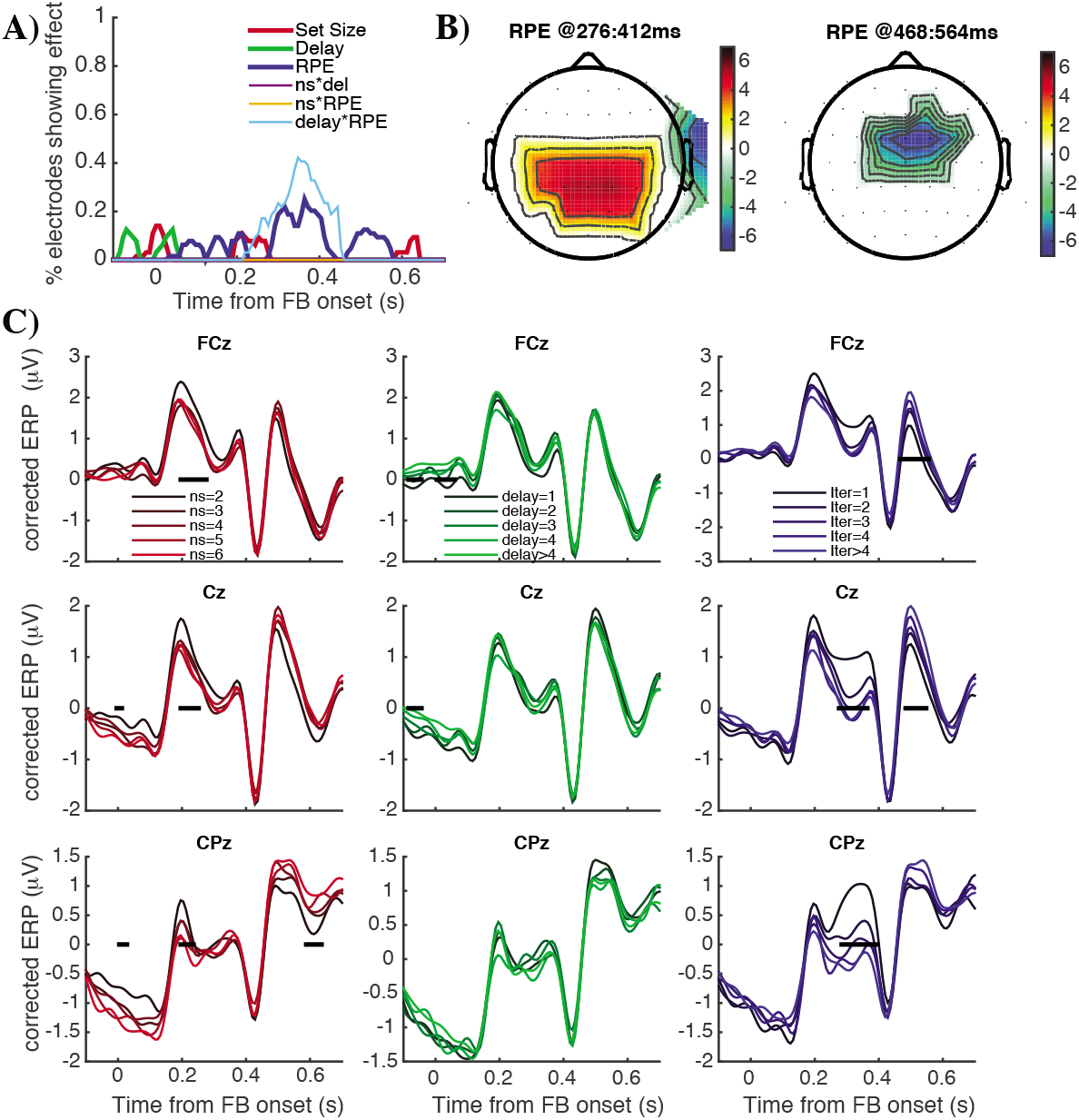
EEG decoding of RL and WM effects during feedback. Feedback-locked effects are plotted with the same conventions as figure 3. RPE: reward prediction error.

### Testing axiomatic indices of RL signals and interactions with WM

We next leveraged these quantities conveying Q-value and RPE signals at distinct time points to test a central axiomatic tenet of neural RL theories (*14*), which, to our knowledge, has not been directly evaluated in neural signal. If neural signals on individual trials truly reflect the latent model variables of expected value and RPE, they should provide more informed estimates of those quantities than those inferred from model fits to behavior alone. Thus, variance in trial-wise stimulus-locked Q value signals should (negatively) predict variance in subsequent feedback-locked RPE signals in the same trial (i.e. greater expectations should be met with diminished surprise), over and above the behavioral RPE (Fig. 5A,B). Indeed, while (by definition) the behavioral RPE accounted for most of the variance in the FB-locked RPE signal (t(38)=10.9, p<10^-4^), increases in trial-wise neural metrics of expected Q value were associated with *lower* neural indices of RPE (t(38)=-2.08, p=0.045), as expected from the computation of *RPE=reward-Q*.

**Figure 5:**
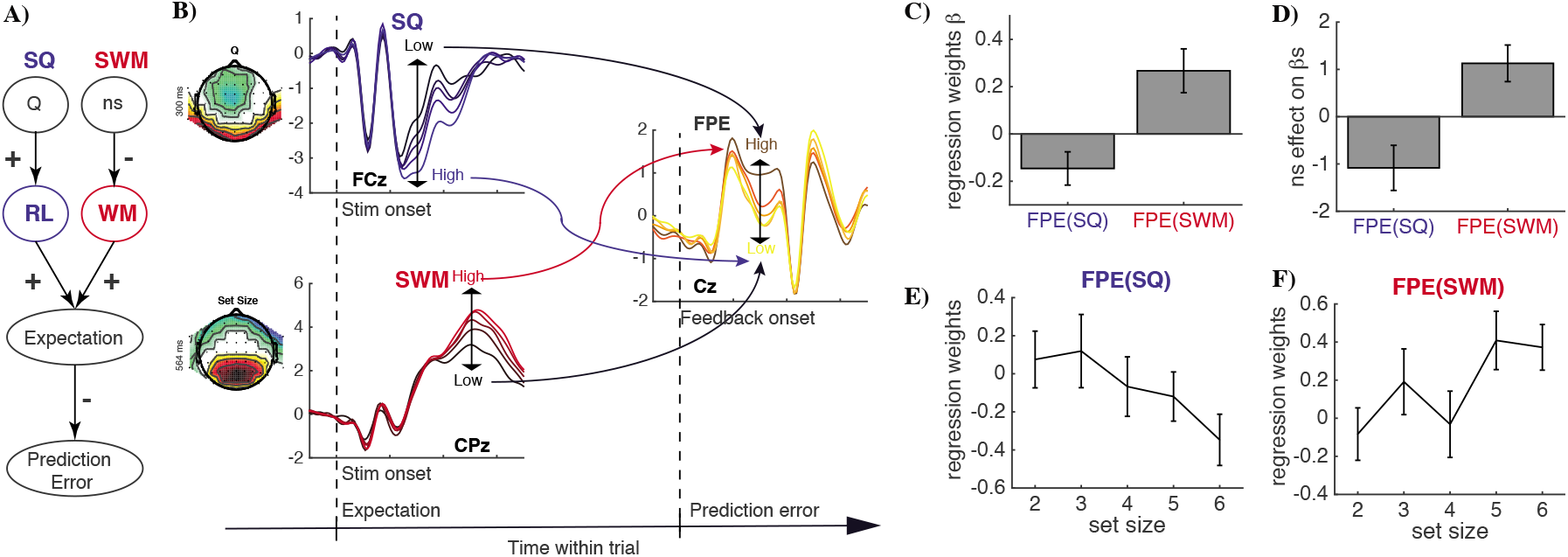
Within-trial dynamics support RL and WM contributions to learning. **A)** A basic tenet of RL is that reward expectation is negatively correlated to the reward prediction error, through the formula RPE = reward – expectation. Expectations can be informed by both RL and WM systems, where RL accumulates reward experience to estimate value *Q* and WM is less reliable with set-size (*ns*), reducing WM contributions to expectation. **B)** Schematic of the trial-by-trial prediction method. For each trial, we compute a stimulus-locked index of RL-related activity (*SQ*) and of WM-related activity (*SWM*), using similarity to spatio-temporal masks obtained from the multiple regression analysis (Fig.3). The model in panel A predicts that trial-by-trial variability in *SQ* negatively predicts neural responses to prediction error during feedback (FPE) of that same trial, with the opposite effect of *SWM*, after controlling for the behavioral reward prediction error. Scalp topographies of SQ and SWM at early and late time points are displayed to the left of each index. **C)** Trial-by-trial variance in FPE is significantly and oppositely accounted for by variability in *SQ* and *SWM*. D-F) These effects are amplified in high set sizes, in which the RL system is relatively more potent (*1–3*).

If the RL and WM processes are independent, neural indices of RL should be independent of set-size. Conversely, if RL processing is degraded when the task becomes more difficult, one might expect that these indices would degrade with set size. Instead, we observed strong evidence for the opposite effect: RL indices were actually enhanced under high load, for both stimulus- and feedback-locked activity (Fig. 6 A t(38)=2.4, p=0.02; Fig. 6B, t(38)=4.5, p=10^-4^). This result is inconsistent with independent RL and WM processes, and instead suggests an interaction between them; but what is the form of this interaction?

**Figure 6:**
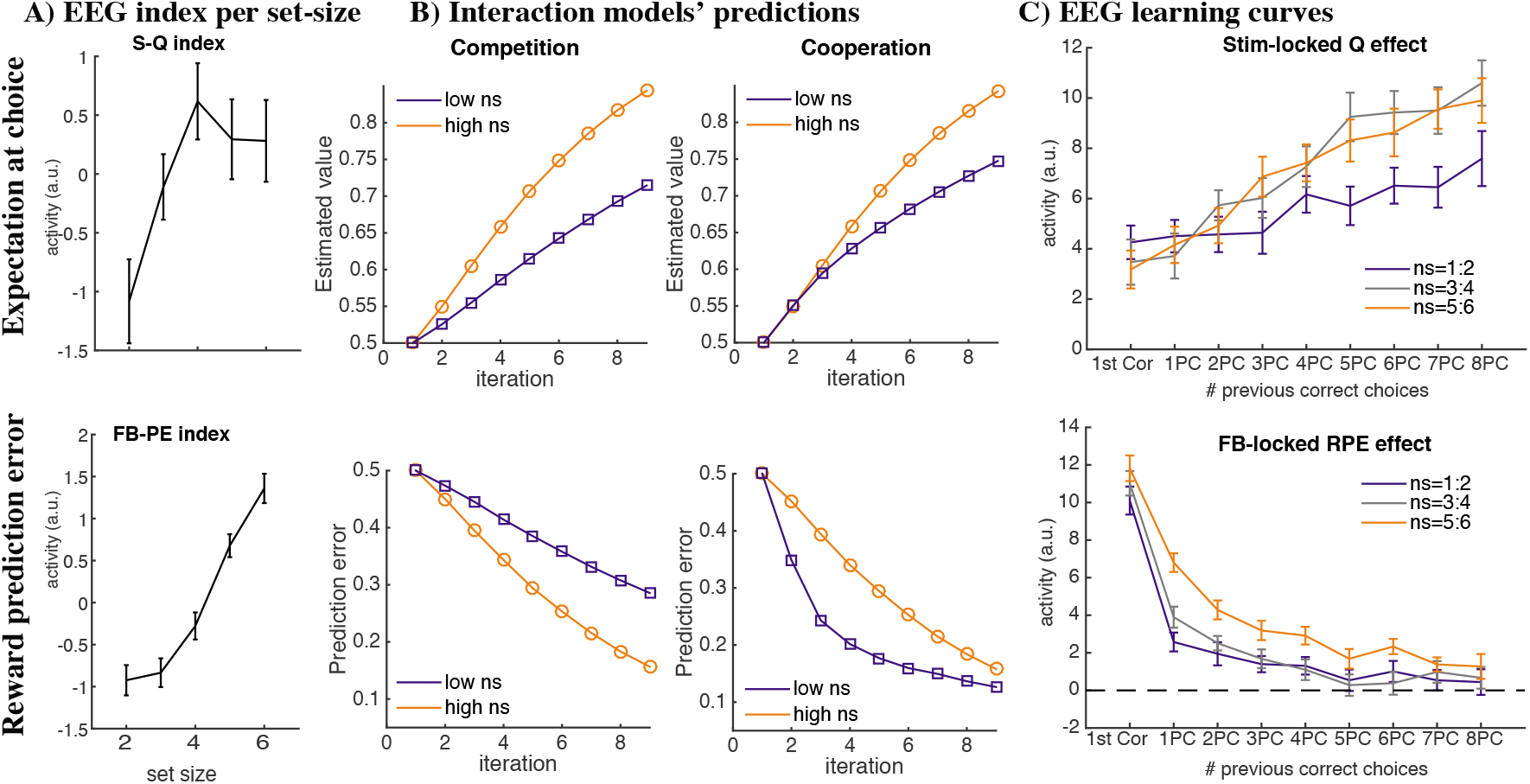
Distinguishing cooperative vs. competitive RL-WM interactions via temporal dynamics of EEG decoding, for stimulus-locked expectation (top) and feedback-locked reward prediction errors (bottom). **A)** Neural markers of RL at both stimulus presentation (*SQ*) and feedback (*FPE*) are amplified with higher set size, suggesting that WM and RL are not independent. **B)** Computational model simulations show that both a competitive and a cooperative account can account for blunted RL computations with higher set sizes (top). However, the competitive account predicts that reward prediction errors decline more rapidly across stimulus iterations in high (orange) compared to low (black) set sizes, whereas the cooperative model predicts the opposite pattern. **C)** Top: stimulus-locked indices of RL increase with learning (by definition), and do so more slowly in low set sizes, consistent with the observed interactions and with both computational models. Bottom: Feedback-locked indices of RPE decrease with learning, and do so more slowly in high set sizes, consistent with the cooperative and refuting the competitive model.

### Distinguishing Competitive from Cooperative Accounts via EEG temporal dynamics

Recent neuroimaging and behavioral findings (*3, 7*) (see also (*15*)) suggested RL-WM interactions during learning, but could not distinguish whether this interaction was cooperative or competitive. Under the competitive hypothesis, the WM system would compete with, and hence hinder, the RL system from learning when WM is reliable (i.e., during low set sizes; Fig. 6B top). Under the cooperative hypothesis, the WM system would instead inform reward expectations needed to compute RPE. Hence, within low set sizes, RPEs would be smaller compared to those expected by pure RL (Fig. 6C top). Thus, both hypotheses can account for the blunted RL signaling observed in low set sizes (Fig. 6A). However, they make qualitatively different predictions regarding the dynamic changes in the RPE signal across trials (Fig. 6B bottom). Specifically, the cooperative account predicts that RPE signals decline more rapidly in low set-sizes (due to the contributions of the fast learning WM system to expectations). In contrast, the competitive model predicts the opposite pattern: when WM dominates in low set sizes, it suppresses learning in the RL system, and hence RL RPEs persist for more trials (compared to high set sizes, where RL learning is unhindered). We tested these hypotheses by plotting the average stimulus- and feedback-locked RL indices as a function of the number of previous correct choices per stimulus, thus obtaining “neural learning curves”. As predicted by both models, learning curves for neural measures of expected Q values were blunted under low load (Fig. 6C top). But supporting the cooperative, but not competitive model, the neural signal of RPEs declined more rapidly under low set sizes than high set-sizes (Fig. 6C bottom; t(38)=3.93, p=0.0003).

To further test this hypothesis, we investigated within-trial temporal dynamics indicative of cooperation. In particular, we asked whether trial-by-trial variance in neural markers of WM during stimulus presentation could predict RPE-related variance in the subsequent feedback period. Specifically, if WM cooperates with RL expectations to compute RPEs, then a high perceived WM load decoded by the stimulus-locked WM index should be predictive of a larger feedback-locked RPE (Fig. 5A). We tested this in a multiple regression model that simultaneously accounted for variance in Q value signals, WM signals, and RPE estimated by fits to behavior alone

Results show that indeed, neural metrics of WM (corresponding to higher perceived load) during stimulus onset were associated with *larger* feedback-locked RPE signals (Fig. 5C; t(38)=2.89, p=0.006). These positive effects contrast with the negative effects of Q value signals (which support prediction error computations), and are predicted by the cooperation model, whereby higher perceived load is indicative of degraded WM representations and hence RPE signals are enhanced relative to when WM is intact. Figs. 5C-F show that these effects hold within individual objective set-sizes, suggesting that the neural index of perceived load is more predictive than objective load, and indeed, are even more pronounced under high set sizes, when WM and RL are more likely to jointly contribute to behavior (SQ: t(38)=-2.27, p=0.03; SWM: t(38)=2.9, p=0.006). Finally, we confirmed the robustness of this double dissociation using a bootstrapping method. Specifically, we shuffled the masks used to obtain trial-level indices of SQ and SWM, and found that the more similar shuffled masks were to Q value mask, the more they predicted decreased neural RPEs, and the more similar they were to WM masks, the more they predicted increased neural RPEs (ps<.0006).

## Discussion

Our findings support a growing literature implicating multiple separable neural processes contributing jointly to human instrumental learning and decision-making (*1*–*4, 7*). While many imaging studies implicate interactive systems with strong debates about how they are arbitrated for choice (*3, 7, 15*–*18*), they have not resolved the specific nature of these interactions, either for choice or for learning. Our model-based decoding of single trial EEG dynamics revealed cooperation between WM and RL systems during learning, despite competition between them for choice.

Specifically, our multiple regression analysis identified two separable spatiotemporal networks sensitive to dissociable aspects of learning. Early on during the choice period, the neural signal was sensitive to reward history, a marker of model-free RL, whereas a later onset signal was sensitive to set size, a marker of working memory load. These results seem to indicate a shift from an early recruitment of a fast, automated RL process to a more deliberative WM (*19*); a conclusion supported by our finding that the early signals were more strongly recruited when WM would be weaker (with increased intervening trials), and thus favoring RL recruitment for decision-making.

Within-trial correlations between choice and feedback dynamics confirmed first, that these are neural signatures of RL, and second, that the RL system is informed by WM for learning. First, trial-by-trial variations of signal encoding *Q*-values during expectations were negatively predictive of variation in that same trial of signals encoding *RPE*, providing evidence for the notion that neural RPE signals compute *reward – Q-value*. These data provide axiomatic evidence (*14*) for a central but here-to-fore untested account of neural RL via within-trial dynamics. Second, in contrast to the neural signature of Q, we found those indicative of higher WM load during expectations were positively related to subsequent RPE signals. These findings provide dissociable signals related to WM and RL expectation that exhibit differential effects on RPE signals as predicted by the cooperation model. Moreover, both of these findings accounted for variance in RPE signals over and above those that could be predicted based on model fits to behavior alone, providing further confirmation that they are related to the computations of interest and evidence that neural markers can be used as a more direct lens into value and decision computations (*20*–*23*).

While much past research has argued for competition between multiple systems during choice (*1, 4, 17*), these studies usually still assume that the systems learn independently, or even compete for learning (*24*). By contrast, our previous behavioral and fMRI findings hinted that the RL computations were not independent of WM, and indeed that the RL process was actually stronger in more difficult task settings, under high WM load (*3, 7*). Our findings here show that indeed, trial-wise EEG markers of the RL process were stronger with increasing WM load, during both decision and feedback periods. Moreover, previous studies could not disentangle cooperative from competitive interactions. The dynamic decoding analysis employed here clearly favors a cooperative mechanism during learning, whereby information held in WM can augment expectations of reward within RL, and thereby reduce subsequent RPEs. These findings directly contrast that predicted by a competitive account of learning, in which reliable WM signals would suppress RPEs within an independent RL system (Fig 6). Second, we showed that trial-by-trial variability in WM signaling in the neural signal at the time of decision predicted variability in the RL signal during subsequent feedback. Together, these findings strongly support our proposed cooperative mechanism, which was not possible in previous studies.

It is interesting to note that in this case, the “cooperative” mechanism actually interferes with the reinforcement learning computation. By decreasing the magnitude of the reward prediction error before the estimate of the Q-value has converged, it slows the learning of the RL Q-values, and thus diminishes RL computations overall. This mechanism predicts that statistical learning of expected reward values would be degraded under low load, a phenomenon we observed behaviorally in a variant of this task using multiple reward outcomes (*7*). This phenomenon can be related to blocking, whereby learning does not occur if another stimulus is predictive of reward (*25*–*27*). Here, however, it is not another stimulus that is predictive of the outcome, but another neural system that more reliably predicts information for the same stimulus (see also (*28*) for an analogous model of hippocampus vs cortex). However, while WM might hinder RL learning in this task, this interaction may be useful in general, allowing WM to be used judiciously for tasks that are less well learned and the RL system to take over when it has accumulated sufficient information. Indeed, since the RL computations occur earlier in the trial, if they are sufficiently reliable, the learner might learn to use only RL and not recruit WM, as we observe occurs over the course of learning (*1, 19*).

To conclude, our results contribute to a better understanding of human learning. First, they show evidence of separable neural processes of WM and RL contributing to learning and competing for decisions in the EEG signal. Second, they provide trial-by-trial evidence for computation of reward prediction errors in the EEG signal related to the RL process. Third, our results provide evidence for a cooperative interplay between WM and RL systems for learning, despite a competitive dynamic during choice. Identifying the neural correlates of the multiple systems that jointly contribute to human learning and decision making is crucial to better understand dysfunction (*2, 7, 29*). Our results are thus an important step toward better understanding of learning in healthy and patient populations.

## Acknowledgements

The authors thank Julie Helmers for help with data collection, as well as Matthew Nassar, Samuel McDougle, Nicholas Franklin, and Andrew Westbrook for comments on this manuscript. This work was funded by grant NSF 1460604 to Michael J Frank and Anne GE Collins.

## Materials and Methods

### 1 Subjects

#### EEG experiment

We collected data for 40 subjects (28 female, ages 18-29), and all were included in the behavioral analyses. 1 subject was excluded from EEG analysis due to technical problems with the EEG cap.

### 2 Experimental Protocol

#### Structure

Subjects performed a learning experiment in which they used reinforcement feedback to figure out which key to press for each presented visual stimulus. The experiment was divided into 22 blocks, with new visual stimuli in each block. After stimulus presentation, subjects selected one of three keys to press with their right hand. Feedback indicated truthfully whether they had selected the correct action for the current stimulus. See “trials” section below for more details.

#### Blocks

Blocks varied in the number of stimuli that participants learned concomitantly (the set size ns) between 1 and 6. Specifically, the number of blocks for set sizes 1-6 were in order {3,6,4,3,3,3}; this number was chosen to ensure at least 12 stimuli and three learning bocks per set size, with the exception of set-size 1, which was used as a control. Within a block, each stimulus was presented a minimum of 9 times and a maximum of 15 times; the block ended after ns*15 trials, or when subjects reached a performance criterion whereby they had selected the correct action for 3 out of the 4 last iterations of each stimulus. Stimulus presentation was pseudorandomized. Stimuli in a given block were all from a single category (e.g. colors, fruits, animals), and did not overlap.

#### Trials

Stimuli were presented centrally on the black background screen (approximate visual angle of 8°); subjects had up to 1.4s to answer by pressing one of three keys with their right hand. Key press was followed by audio-visual feedback presentation (word “Win!”, ascending tone, or “loss”, descending tone), with a uniformly jittered lag of .1-.6s. Failure to answer within 1.4s was indicated by a “too slow” message. Feedback was presented for [.4-.8]s, and followed by a [.5-.8]s fixation cross before next trial onset.

### 3 Model free analysis

We analyze behavior using a multiple logistic regression. Predictors include set size, delay (number of trials since last previous correct choice for current trial’s stimulus), iterations (number of previous correct trials for current trial’s stimulus), and interactions between those factors. The first two predictors are markers of WM function, and are also used for the EEG multiple regression analysis. The third regressor is a marker of reward history and thus targets the RL system. Following previous published methods, main effect predictors were transformed according to X → -1/X.

### 4 Computational modeling

#### RLWM model

To better account for subjects’ behavior and disentangle roles of working memory and reinforcement learning, we fitted subjects’ choices with our hybrid RLWM computational model. Previous research showed that this model, allowing choice to be a mixture between a classic delta rule reinforcement learning process and a fast but capacity-limited and delay-sensitive working memory process, provided a better quantitative fit to learning data than models of either WM or RL alone (*1, 30*). The model used here is identical to the model used in (*3*). We first summarize its key properties, following by the details:

- RLWM includes two modules which separately learn the value of stimulus-response mappings: a standard incremental procedural RL module with learning rate α, and a WM module that updates S-R-O associations in a single trial (learning rate 1) but is capacity-limited (with capacity K).
- The final action choice is determined as a weighted average over the two modules’ policies. How much weight is given to WM relative to RL (the mixture parameter) is dynamic and reflects the probability that a subject would use WM vs. RL in guiding their choice. This weight depends on two factors. First, a constraint factor reflects the a priori probability that the item is stored in WM, which depends on set size n_S_ of the current block relative to capacity K (i.e., if n_S_>K, the probability that an item is stored is K/ns), scaled by the subject’s overall reliance of WM vs. RL (factor 0<ρ<1), with higher values reflecting relative greater confidence in WM function. Thus, the constraint factors indicates that the maximal use of WM policy relative to RL policy is w_0_ = ρ⨯ min(1, K/n_S_). Second, a strategic factor reflects the inferred reliability of the WM compared to RL modules over time: initially, the WM module is more successful at predicting outcomes than the RL module, but because it has higher capacity and less vulnerability to delay, the RL module becomes more reliable with experience.
- Both RL and WM modules are subject to forgetting (decay parameters ϕ_RL_ and ϕ_WM_). We constrain ϕ_RL_ < ϕ_WM_ consistent with WM’s dependence on active memory).

### Learning model details

#### Reinforcement learning model

All models include a standard RL module with simple delta rule learning. For each stimulus *s*, and action *a*, the expected reward *Q(s,a)* is learned as a function of reinforcement history. Specifically, the *Q* value for the selected action given the stimulus is updated upon observing each trial’s reward outcome r_t_ (1 for correct, 0 for incorrect) as a function of the prediction error between expected and observed reward at trial *t*:

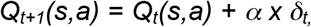

where δ_*t*_= *r_t_* - *Q_t_*(*s*,*a*) is the prediction error, and *α* is the learning rate. Choices are generated probabilistically with greater likelihood of selecting actions that have higher *Q* values, using the softmax choice rule:

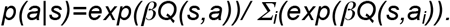

Here, *β* is an inverse temperature determining the degree with which differences in *Q*-values are translated into more deterministic choice, and the sum is over the three possible actions a_i_. Because we have found that within this experimental protocol, recovering *β* independently from the learning rate is often impractical, we fix *β*=100.

#### Undirected noise

The softmax temperature allows for stochasticity in choice, but where stochasticity is more impactful when the value of actions are similar to each other. We also allow for “slips” of action (“irreducible noise”, i.e., even when Q value differences are large). Given a model’s policy π = p(a|s), adding undirected noise consists in defining the new mixture policy:

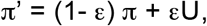

where U is the uniform random policy (U(a) = 1/n_A_, n_A_=3), and the parameter 0<ε<1 controls the amount of noise (*31–33*). (*34*) showed that failing to take into account this irreducible noise can render fits to be unduly influenced by rare odd data points (e.g. that might arise from attentional lapses), and that this problem is remedied by using a hybrid softmax-ε-greedy choice function as used here.

#### Forgetting

We allow for potential decay or forgetting in Q-values on each trial, additionally updating all Q-values at each trial, according to:

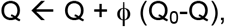

where 0<ϕ<1 is a decay parameter pulling at each trial the estimates of values towards initial value Q_0_ = 1/n_A_. This parameter allows us to capture delay-sensitive aspects of WM, where active maintenance is increasingly likely to fail with intervening time and other stimuli, but also allows us to separately estimate any decay in RL values (which is typically substantially lower than in WM).

#### Perseveration

To allow for potential neglect of negative, as opposed to positive feedback, we estimate a perseveration parameter *pers* such that for negative prediction errors (delta<0), the learning rate α is reduced by α = (1-*pers*) x α. Thus, values of *pers* near 1 indicate perseveration with complete neglect of negative feedback, whereas values near 0 indicate equal learning from negative and positive feedback.

#### Working Memory

To implement an approximation of a rapid updating but capacity-limited WM, this module assumes a learning rate α = 1 (representing the immediate accessibility of items in active memory), but includes capacity limitation such that only at most K stimuli can be remembered. At any trial, the probability of working memory contributing to the choice for a given stimulus is *w*_*WM*_*(t) =P*_*t*_*(WM)*. This value is dynamic as a function of experience (see next paragraph). As such, the overall policy is:

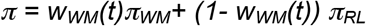

where πWM is the WM softmax policy, and πRL is the RL policy. Note that this implementation assumes that information stored for each stimulus in working memory pertains to action-outcome associations. Furthermore, this implementation is an approximation of a capacity/resource-limited notion of working memory. It captures key aspects of working memory such as 1) rapid and accurate encoding of information when low amount of information is to be stored; 2) decrease in the likelihood of storing or maintaining items when more information is presented or when distractors are presented during the maintenance period; 3) decay due to forgetting. Because it is a probabilistic model of WM, it cannot capture specifically which items are stored, but it can provide the likelihood of any item being accessible during choice given the task structure and recent history (set size, delay, etc).

#### Inference

The weighting of whether to rely more on WM vs. RL is dynamically adjusted over trials within a block based on which module is more likely to predict correct outcomes. The initial probability of using WM *w*_*WM*_*(0) = P*_*0*_*(WM)* is initialized by the *a priori* use of WM, as defined above, *w*_*WM*_*(0) = ρ x min(1, K/n*_*S*_), where *ρ* is a free parameter representing the participant’s overall reliance on WM over RL.

On each correct trial, *w*_*WM*_*(t)=P*_*t*_*(WM)* is updated based on the relative likelihood that each module would have predicted the observed outcome given the selected correct action a_c_; specifically:

- for WM, p(correct|stim, WM) = wWM π WM(ac) + (1-w WM)1/nA
- for RL, p(correct|stim, RL) this is simply π RL(ac)

The mixture weight is updated by computing the posterior using the previous trial’s prior, and the above likelihoods, such that

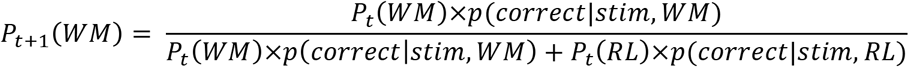

and *P_t+1_*(*RL*)=*1*-*P_t+1_*(*WM*).

#### Models Considered

We combined the previously described features into different learning models and conducted extensive comparisons of multiple models to determine which fit the data best (penalizing for complexity) so as to validate the use of this model in interpreting subjects’ data. For all models we considered, adding undirected noise, forgetting and perseveration features significantly improved the fit, accounting for added model complexity (see model comparisons).

This left three relevant classes of models to consider:

- **RLF**: This model combines the basic delta rule RL with forgetting, perseveration and undirected noise features. It assumes a single system that is sensitive to delay and asymmetry in feedback processing. This is a 4-parameter model (learning rate α, undirected noise ε, decay ϕRL, and *pers* parameter).
- **RL6**: This model is identical to the previous one, with the variant that learning rate can vary as a function of set-size. We have previously shown that while such a model can capture the basic differences in learning curves across set-sizes by fiting lower learning rates with higher set sizes, it provides no mechanism that would explain these effects, and still cannot capture other more nuanced effects (e.g. changes in the sensitivity to delay with experience). However it provides a benchmark to compare with RLWM. This is a 9-parameter model (6 learning rate αns, undirected noise ε, decay ϕRL, and *pers* parameter).
- **RLWM**: This is the main model, consisting of a hybrid between RL and WM. RL and WM modules have shared *pers* parameter, but separate decay parameters, ϕRL and ϕWM, to capture their differential sensitivity to delay. Working memory capacity is 0<K<6, with an additional parameter for overall reliance on working memory 0<ρ<1. Undirected noise is added to the RLWM mixture policy. This is an 8-parameter model (capacity K, WM reliance ρ, WM decay ϕWM, RL learning rate α, RL decay ϕRL, undirected noise ε, and *pers* parameter).

In the RLWM model presented here, the RL and WM modules are independent, and only compete for choice at the policy level. Given our findings showing an interaction between the two processes, we also considered variants of RLWM including mechanisms for interactions between the two processes at the learning stage. These models provided similar fit (measured by AIC) to the simpler RLWM model. We chose to use the simpler RLWM model, because the more complex model is less identifiable within this experimental design, providing less reliable parameter estimates and regressors for model-based analysis.

### RLWM fitting procedure

We used matlab optimization under constraint function fmincon to fit parameters. This was iterated with 50 randomly chosen starting points, to increase likelihood of finding a global rather than local optimum. For models including the discrete capacity *K* parameter, this fitting was performed iteratively for capacities K = {1,2,3,4,5}, using the value gave the best fit in combination with other parameters. All other parameters were fit with constraints [0 1].

### Model Comparison

We used the Akaike Information Criterion to penalize model complexity - AIC (*35*). Indeed, we previously showed that in the case of the RLWM model and its variants, AIC was a better approximation than Bayesian Information Criterion (BIC; Schwarz 1978) at recovering the true model from generative simulations (*1*). Comparing RLWM, RL6 and RLF showed that models RL6 and RLF were strongly non-favored, with exceedance probability for RLWM of 0.95 over the whole group (*37*). Other single process models were also unable to capture behavior better than RLWM.

### Model Simulation

Model selection alone is insufficient to assess whether the best fitting model sufficiently captures the data. To test whether models capture the key features of the behavior (e.g., learning curves), we simulated each model with fit parameters for each subject, with 100 repetitions per subject then averaged to represent this subject’s contribution. In order to account for initial biases, we assume that the model’s choice at first encounter of a stimulus is identical to the subjects, while all further choices are randomly selected from the model’s learned values and policies.

## 5 Interaction models

We test two computational models embodying two distinct hypotheses for WM and RL interactions, that both predict the low set-size blunted RL observed experimentally.

The *competitive* model assumes that in low set sizes where WM is successful, it inhibits RL computations, such that the prediction error is computed normally δ=*R–Q_RL_*, but the update is weakened *Q_RL_=Q_RL_+*αηδ, where η indicates the degree of interference of WM in the RL computation.

The *cooperative* model instead assumes that in low set sizes where WM is successful, it contributes part of the reward expectation for the RL model, according to the equation: δ = *R – [*η*Q_RL_ + (1+*η*)Q_WM_]*, where *Q*_*WM*_ represents reward expectations from the working memory system. This reward prediction error is then used to update RL as normal: *Q_RL_= Q_RL_+*αδ. Because WM learns quickly, WM contribution makes d smaller than expected from classic RL, and thus leads to blunted RL.

Simulations for fig. 6 were run with the following parameters for both models:α=.2, and η=0 or .5 for high or low working memory, respectively.

We did not fit the behavior with the interaction models because this experimental design is not appropriate to capture behavioral markers of interaction (contrary to (*7*)), and thus these models’ parameters are not satisfactorily identifiable. Assuming independence between RL and WM in the model-fitting allows us to capture behavior well on average (Fig. 2) but also to investigate the degree to which neural signals deviate from independence in the way predicted by the cooperation vs. competition models without assuming either.

## 6 EEG

### 6.1 System

EEG was recorded from a 64-channel Synamps2 system (0.1-100 Hz bandpass; 500Hz sampling rate).

### 6.2 Data Preprocessing/Cleaning

EEG was recorded continuously with hardware filters set from 0.1 to 100 Hz, a sampling rate of 500 Hz, and an online vertex reference. Continuous EEG was epoched around the feedback onset (-1500 to 2500 ms). We used previously identified data cleaning and preprocessing method (*38, 39*) facilitated by the EEGlab toolbox (*40*): data was visually inspected to identify bad channels to be interpolated and bad epochs to be rejected. Blinks were removed using independent component analysis from EEGLab. The electrodes were referenced to the average across channels.

### 6.3 ERPs

For event-related potentials (ERP) and multiple regression analysis, data were bandpass filtered from 0.5 to 20 Hz, down-sampled to 125Hz, and baselined by the mean activity between -100ms and 0ms prior to stimulus onset. For each subject, we performed a multiple regression at each electrode and time point within -100:700ms around stimulus onset (101 time points) and feedback onset. Because there were many fewer error than correct trials, we included only correct trials in the analysis. Scalp voltage was z-scored before being entered into the multiple regression analysis.

In the stimulus-locked analysis, regressors of interest included z-scored set size, delay, model-derived RL expected value, and the interaction of those three regressors; regressors of no interest included reaction time (z-scored log-transformed), and z-scored trial number within block.

Further stimulus-locked analysis mostly focused on regression weights for the main three regressors, β_S-NS_ and β_S-Delay_, considered as markers of WM function, β_S-Q_, marker of RL function; which we obtained for each subject, time-point, and electrode. The feedback locked analysis was identical, but with RL reward prediction error replacing RL expected value, producing key regression weights β_F-NS_ and β_F-Delay_, and β_F-RPE_. Trials included in the analysis were all correct trials for which the values of the regressors were well defined, namely trials of set size 2 and above, with at least one previous correct choice for the current stimulus (insuring delay is defined).

### 6.4 Statistical analysis of GLM weights

we tested the significance of regression weights against 0 across subjects for all electrodes and time-points. To correct for multiple comparisons, we performed cluster-mass correction by permutation testing with custom-written matlab scripts, following the method described (*41*). Cluster formation threshold was for a ttest significance level of p=0.001. Cluster mass was computed across spacetime, and only clusters with greater mass than maximum cluster mass obtained with 95% chance permutations were considered significant, with 1000 random permutations.

### 6.5 Corrected ERPs

To plot corrected ERPs, we compute the predicted voltage by the multiple regression model when setting a single regressor to 0 (set-size, delay, RPE, or reaction time; we the substract this form the true voltage, leaving only the fixed effect, the variance explained by that regressor, and the remaining noise of the regression model.

### 5.6 Trial-by-trial markers of WM and RL

We used sensitivity to set-size as a marker of WM-dependent processing, and sensitivity to model-inferred RL-Q value or RL-RPE as a marker of RL processing. To compute trial-by-trial markers of each process, we first define a spatio-temporal mask as a result of the analysis of GLM weights: the stimulus-locked working memory mask is defined at each time point and electrode by 0 if the effect of set-size is not significant, the t-value of the effect if it is significant. A similar process defines the stimulus-locked Q mask, and the feedback locked RPE mask. Index of WM activation ***SWM*** in a given trial is then computed by how well its activity pattern matches the mask, which we compute as a cross product of the mask by the activity pattern. The same process is applied to stimulus-locked Q-mask, yielding a trial-by-trial index stimulus-locked RL activity ***SQ***, and to feedback-locked RPE-mask, yielding a trial-by-trial index of feedback locked RL activity ***FPE***. EEG learning curves plotted in Fig. 6 show averaged indices, as a function of trial iteration number. We test the model predictions by computing the difference in index from first to second correct trial, and comparing this value between low and high set sizes.

### 6.7 Link between stim-locked and feedback locked

We used the previously defined indices to ask whether working memory and RL related activity at stimulus presentation predicted RL related activity at feedback. To do so, we tried to explain FPE index in a multiple regression including the behavioral reward prediction error and the EEG SWM and SQ indices.

**Figure S1:**
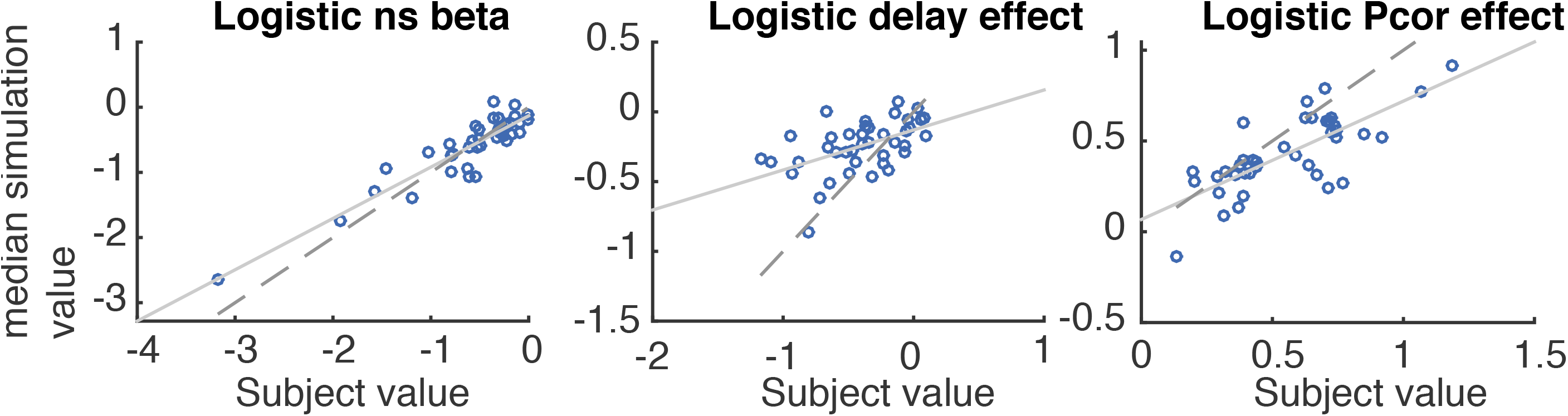
Model validation: logistic regression analysis of model behavior simulated with model parameters fit on participants’ behavior (100 simulations per subject). Median regression weights per subject for set size (ns), delay, and iterations are strongly related to actual participants’ regression weights.

**Figure S2:**
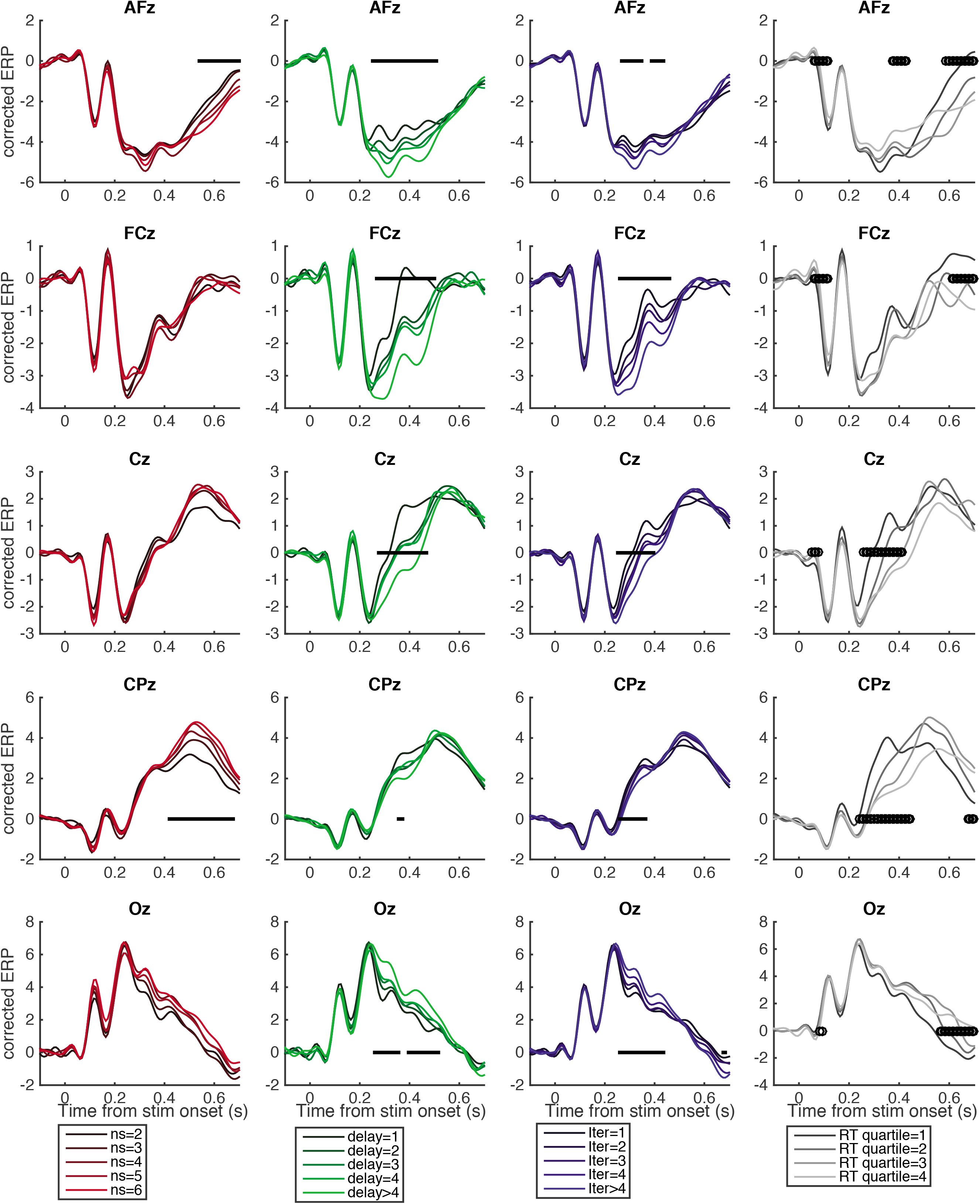
See figure 3B. Right-most column shows corrected ERPs for different reaction time quartiles.

**Figure S3:**
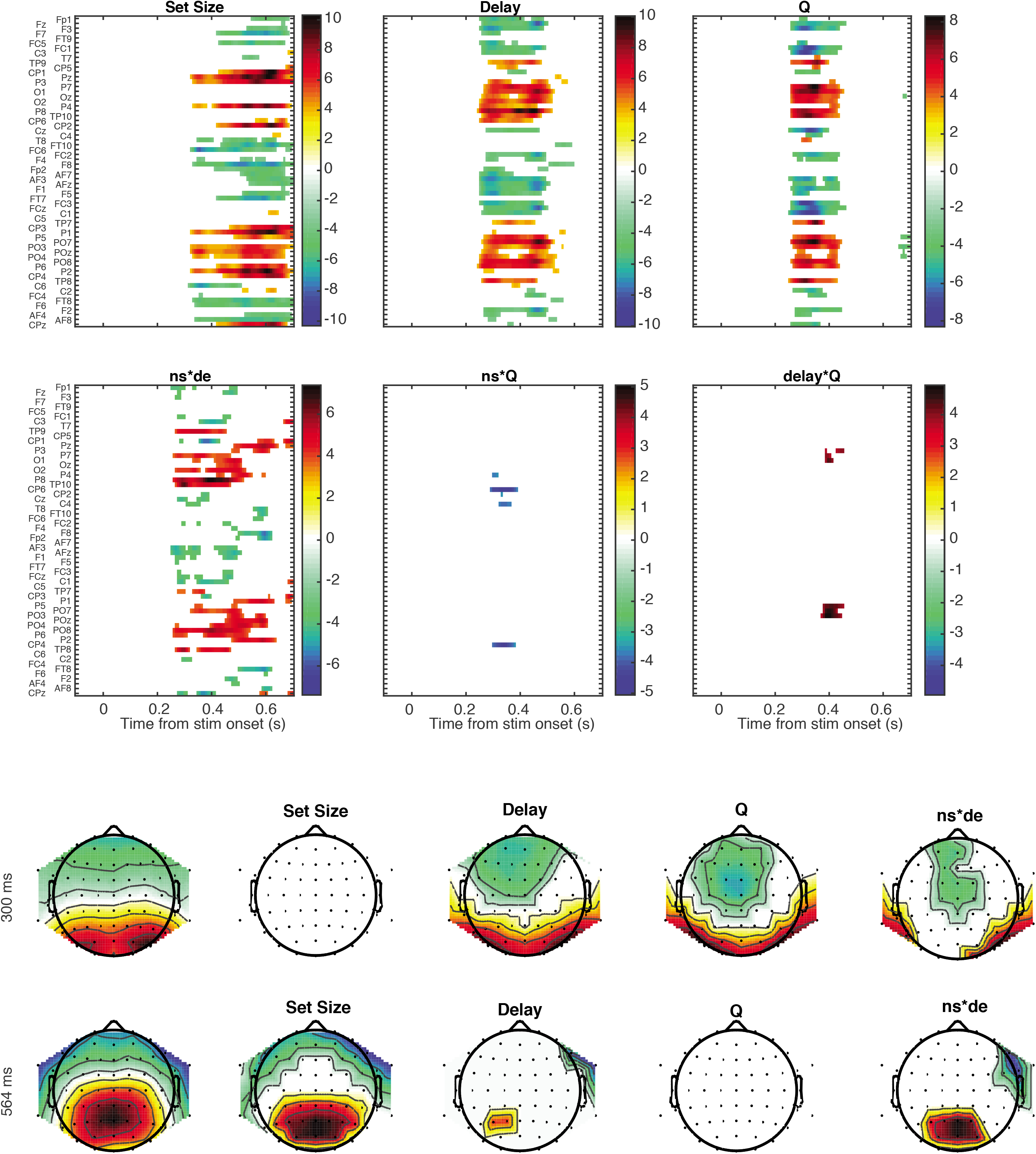
See figure 3C. Top heat-maps show stimulus-locked multiple regression analysis results for all time points and electrodes, as well as for the interactions of the three main factors (set size ns, delay, Q-values).

**Figure S4:**
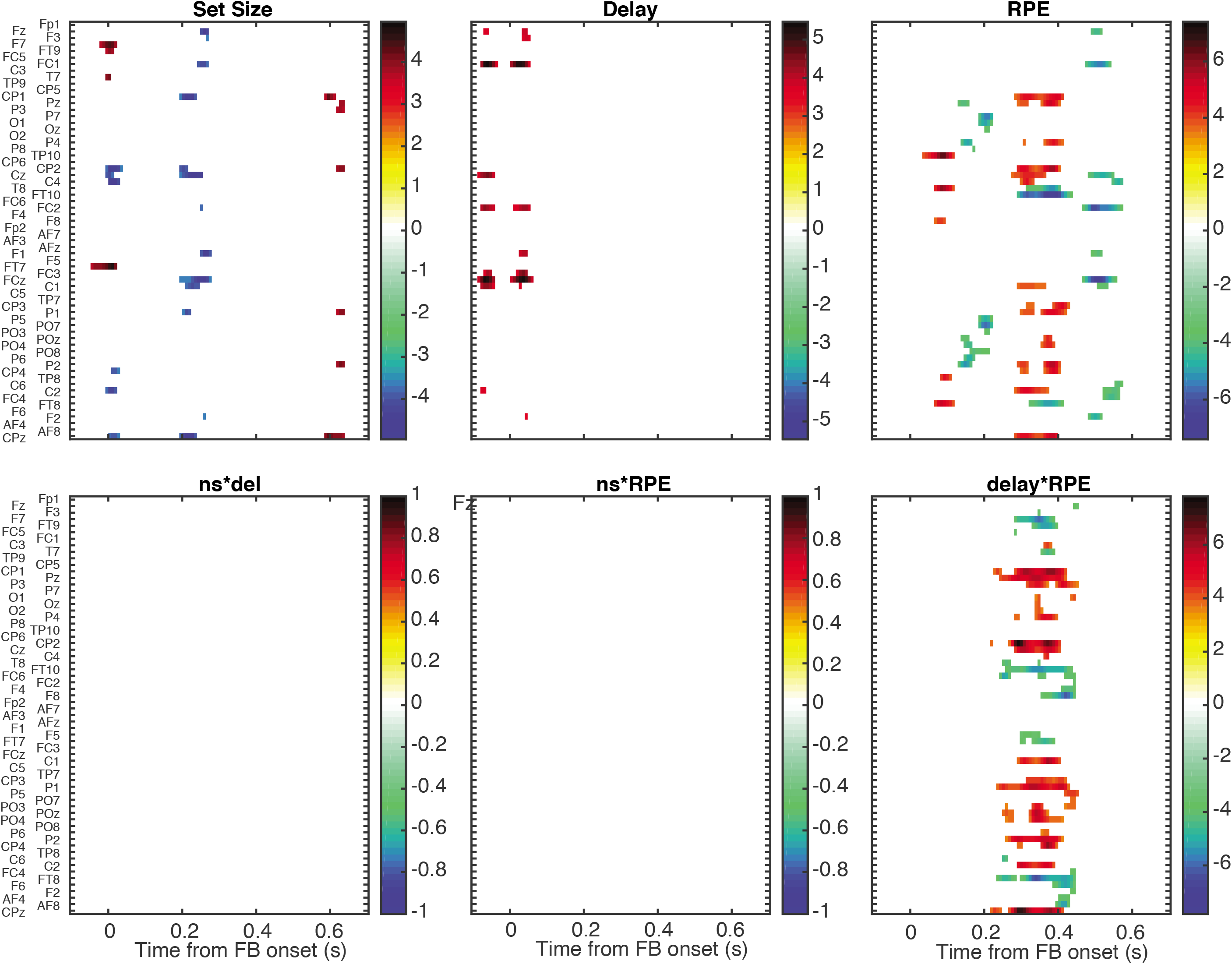
See figure 4; heat-maps show feedback-locked multiple regression analysis results for all time points and electrodes, as well as for the interactions of the three main factors (set size ns, delay, reward prediction errors RPE).

